# Critical interactions for SARS-CoV-2 spike protein binding to ACE2 identified by machine learning

**DOI:** 10.1101/2021.03.19.436231

**Authors:** Anna Pavlova, Zijian Zhang, Atanu Acharya, Diane L. Lynch, Yui Tik Pang, Zhongyu Mou, Jerry M. Parks, Chris Chipot, James C. Gumbart

## Abstract

Both SARS-CoV and SARS-CoV-2 bind to the human ACE2 receptor. Based on high-resolution structures, the two viruses bind in practically identical conformations, although several residues of the receptor-binding domain (RBD) differ between them. Here we have used molecular dynamics (MD) simulations, machine learning (ML), and free energy perturbation (FEP) calculations to elucidate the differences in RBD binding by the two viruses. Although only subtle differences were observed from the initial MD simulations of the two RBD-ACE2 complexes, ML identified the individual residues with the most distinctive ACE2 interactions, many of which have been highlighted in previous experimental studies. FEP calculations quantified the corresponding differences in binding free energies to ACE2, and examination of MD trajectories provided structural explanations for these differences. Lastly, the energetics of emerging SARS-CoV-2 mutations were studied, showing that the affinity of the RBD for ACE2 is increased by N501Y and E484K mutations but is slightly decreased by K417N.

## Introduction

The COVID-19 pandemic, caused by the SARS-CoV-2 Betacoronavirus, remains a global crisis with extensive negative economic, social, and health effects. This is the third emergence of a coronavirus outbreak in the 21st century, with SARS-CoV and MERS-CoV appearing in 2002 and 2012, respectively.^1,2^ Although all three of these viruses belong to the Betacoronavirus genus, SARS-CoV-2 appears to have a higher rate of transmission,^1–3^ and, in contrast to these earlier outbreaks, SARS-CoV-2 has spread rapidly worldwide. Continued transmission promotes the appearance of new mutant strains of the virus. Given both the likelihood of continued SARS-CoV-2 mutations,^4^,^5^ including the recent U.K., South African, and South American variants,^6^ as well as the potential for additional coronavirus zoonotic jumps,^1,7^ there is an urgent need to develop effective strategies to combat such outbreaks. A detailed understanding of the mechanism of viral infection, which begins with virus-host cell recognition, is therefore required to devise these strategies.

Early studies suggested several molecular level factors that contribute to increased infectivity of SARS-CoV-2 relative to SARS-CoV. These coronaviruses utilize a similar mechanism for host cell entry and the initial infection depends critically upon host cell recognition, which is mediated by the viral S (spike) protein^3,8–10^ via the human angiotensin-converting enzyme 2 (ACE2),^9,11–15^ a cell surface protein found in various tissues.^16^ As this proteinprotein interaction initiates infection, the spike protein is an attractive therapeutic target.^17,18^ The spike protein is a homotrimeric type I fusion glycoprotein, which protrudes from the surface of the virus and is composed of two subunits, S1 and S2 (Fig. 1a). While S2 contains the membrane fusion machinery, the S1 region mediates initial recognition and binding to the host cell receptor via its receptor-binding domain (RBD).^8–10^

**Figure 1.**
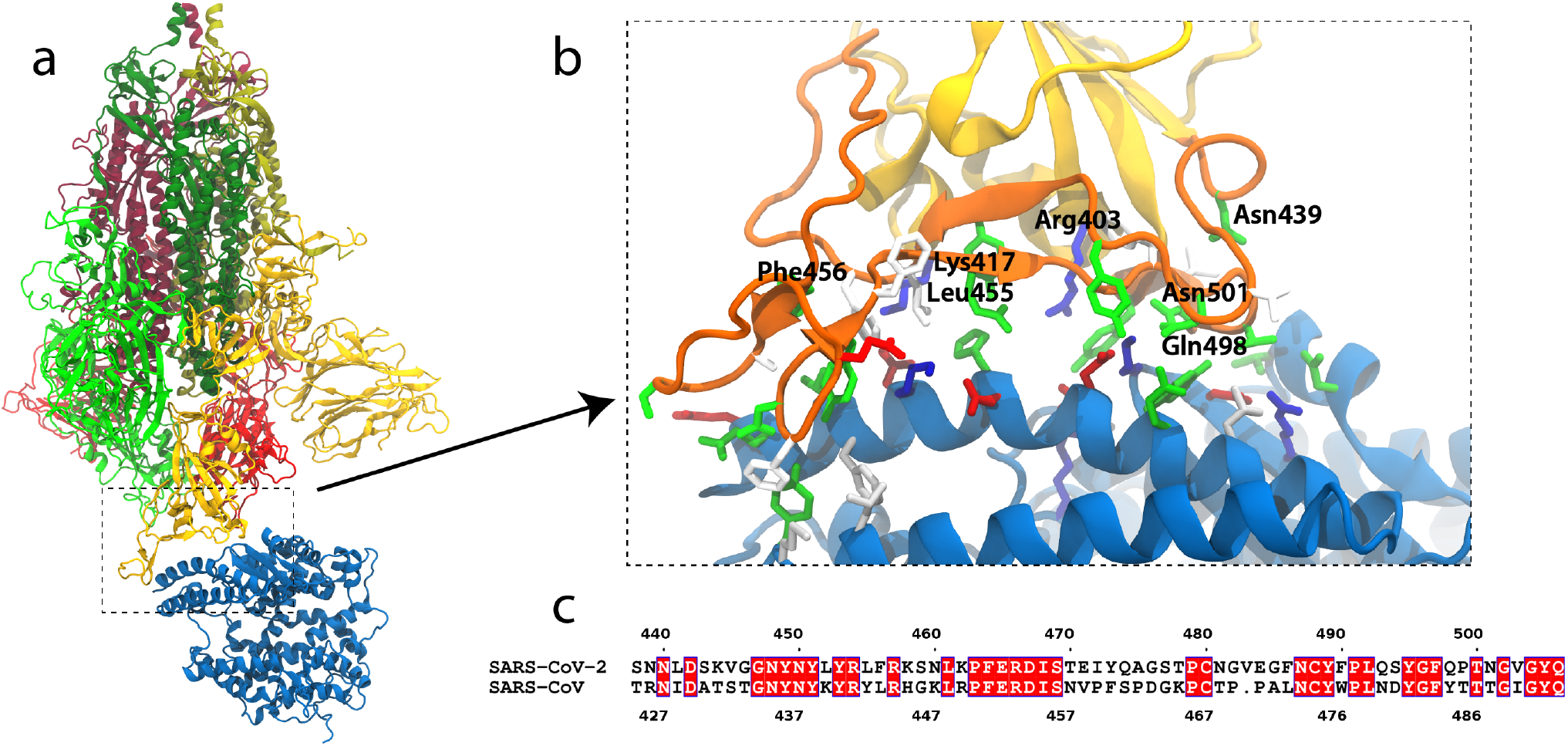
SARS-CoV-2 S-protein binding to ACE2. (a) S-protein trimer (green, yellow, red) bound to the peptidase domain of ACE2 (blue). The lighter and darker shades of the trimer are S1 and S2, respectively. The viral envelope would be at the top and the host cell membrane at the bottom. (b) Close-up of the RBM (orange) interacting with ACE2. Residues near the interface are shown in a stick representation, colored by residue type (blue and red are positively and negatively charged, respectively, green is polar, white is hydrophobic). Specific residues in SARS-CoV-2 are labeled. (c) Alignment of the RBMs of SARS-CoV-2 and SARS-CoV^19^. Identical residues are white on red background.

**Figure 2.**
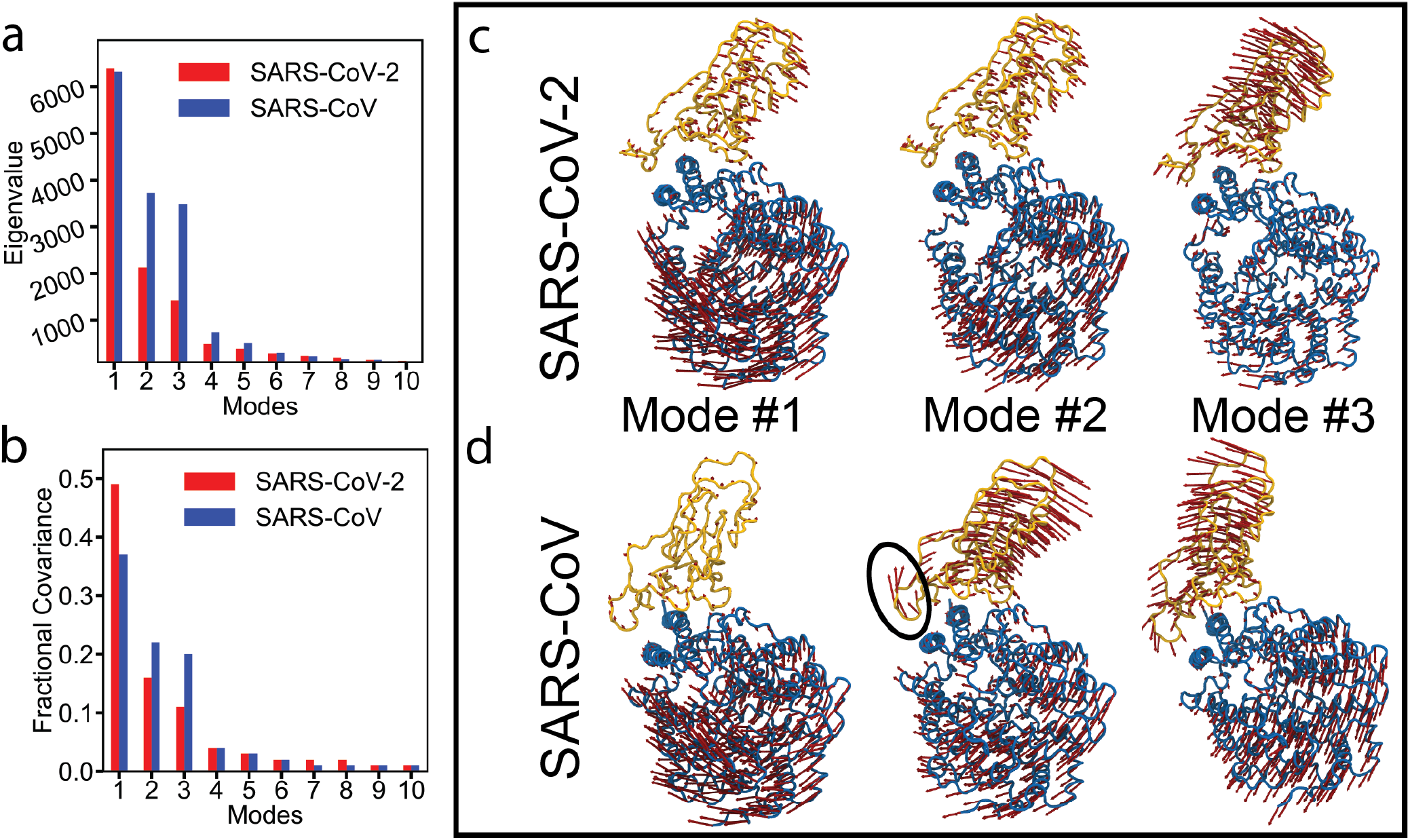
Essential dynamics analysis of two RBD-ACE2 complexes of SARS-CoV-2 and SARS-CoV viruses. (a) Eigenvalues and (b) fractional variance of top-10 modes. Three dominant modes of the complexes with (c) SARS-CoV-2 and (d) SARS-CoV. ACE2 and RBD are shown in blue and yellow, respectively. Motion of a loop in SARS-CoV in mode #2 is highlighted with a black oval.

Although the RBD sequences for SARS-CoV and SARS-CoV-2 are ~74% identical overall, the sequence identity drops to only ~50% for the receptor-binding motif (RBM), the region that contacts ACE2 (Figs. 1b,c).^20,21^ In fact, several recent studies^13^–^15^ have demonstrated that the isolated RBD of SARS-CoV-2 has a slightly higher binding affinity for ACE2 than the corresponding RBD of SARS-CoV, with this enhanced affinity contributing to the increased transmission/infectivity of SARS-CoV-2 relative to SARS-CoV.^3,10^ For SARS-CoV, Wu et al.^22^ have found a correlation between optimized RBD-ACE2 binding and viral entry/infection. Moreover, recent crystal^13,15^ and cryo-EM^12^ structures of the SARS-CoV-2 RBD-ACE2 complex have revealed atomistic details of the binding interface, thereby providing a structural basis to begin to understand the binding affinity differences. A total of 17 residues of SARS-CoV-2 and 16 residues of SARS-CoV were identified to be within 4Å of ACE2, all part of their respective RBMs except Lys417 of SARS-CoV-2, and a majority being identical or at least similar^13^. These sequence differences are spread across the entire binding interface, and how they, as well as recent RBD mutations, collectively contribute to enhanced stability and binding affinity of SARS-CoV-2 for ACE2 is not completely understood.

A number of previous molecular dynamics (MD) studies have compared the dynamics of SARS-CoV and SARS-CoV-2 complexes with ACE2,^21,23–26^ including a study of the full-length complex in the context of the cellular membrane.^27^ It was shown that the SARS-CoV-2 complex is more stable, as measured by root-mean-square deviation (RMSD) and the increased number contacts between RBD and ACE2.^21,24^ In addition, differences in dynamics were observed between the two complexes and binding contributions of several mutations were quantified. Coarse-grained simulations were used to estimate the dissociation constant and revealed that the RBD-ACE2 binding affinity in SARS-CoV-2 is twice that of SARS-CoV.^26^ Furthermore, all-atom steered MD simulations showed that slightly greater force is required to pull the RBD and ACE2 apart in SARS-CoV-2.^26^ In addition, FEP calculations have been performed to assess several amino-acid replacements between SARS-CoV and SARS-CoV-2.^24,28^ Recent studies also highlighted the role of glycans on SARS-CoV-2 spike proteins.^29,30^ While some glycans directly influence the opening or closing of the RBD,^29^ viral glycans do not form direct interactions with ACE2.^30^

Here, we employ multiple-μs MD simulations to compare the relative stabilities, dynamics, and energetic differences between the SARS-CoV RBD-ACE2 and SARS-CoV-2 RBD-ACE2 complexes. Recent advances in both hardware and software^31,32^ have allowed MD simulations to be performed on increasingly large systems over long timescales. These simulations produce an extraordinary amount of data, which poses serious challenges for analysis and interpretation.^33–38^ Similar to the recent work of Fleetwood et al.,^34^ we use supervised machine learning (ML) approaches to assist in identifying those residues that contribute the most to dynamical differences between the two viral RBDs. Based on this identification, we further quantify the relative changes in binding free energy for the RBD with ACE2 via free energy perturbation (FEP) calculations. Our approach reveals subtle distinguishing residue-level differences between the complexes with FEP results that are in accordance with available experimental data. Although we have focused on the SARS-CoV and SARS-CoV-2 complexes with ACE2, the combination of MD, ML, and FEP provides an attractive approach for understanding essential differences at the atomic level and is expected to be useful for other systems as well.

## Results

### Global interactions between RBD and ACE2

To discriminate between the binding of the SARS-CoV and SARS-CoV-2 RBDs to ACE2, we ran two 2-μs simulations, starting each from the structurally characterized bound states (PDBs 2AJF and 6M17, respectively; see Methods). We monitored the RMSD of the RBD and ACE2 individually in each simulation. The RMSD of SARS-CoV RBD from both runs quickly rises above those of SARS-CoV-2 RBD (Fig. S1a), due in part to the loss of a small helix near the N-terminus in the former (residues 326-330). Similarly, the RMSD of ACE2 in both SARS-CoV runs rises above those from the SARS-CoV-2 runs (Fig. S1b), in line with previous studies.^21,24,39^

We also measured the root-mean-square fluctuations (RMSF) for each trajectory (Fig. S1e). One of the largest differences is seen in the RBD loop containing residues 380-400 (367-387 in SARS-CoV), which is more dynamic in SARS-CoV than in SARS-CoV-2. This loop is on the side of the RBD opposite to the interface with ACE2 and includes residues missing from the crystal structure (376-381).^11^ Similarly, the loop containing residues 453-473 (440-460 in SARS-CoV) in the RBM is consistently more dynamic in SARS-CoV compared to SARS-CoV-2. In contrast, in one simulation of the SARS-CoV-2 complex, the loop containing residues 442-451 (429-438 in SARS-CoV) is destabilized, moving away from the interface.

The bound state was maintained in all simulations. However, the loop containing residues 458-475 (SARS-CoV numbering) was observed to move away from ACE2 twice in the SARS-CoV simulations (Fig. S2). This movement was not observed for SARS-CoV-2, suggesting a tighter binding, in agreement with previous experimental data.^14,15^ Although the hydrogen-bond count remained the same in both simulations, this motion resulted in a temporary reduction in the number of hydrophobic contacts for SARS-CoV (Figs. S1c,d). The different behavior observed for this loop is in line with the large number of differences in sequence in this region of the viral RBM (Fig. 1c).

We also performed essential dynamics analysis (EDA)^40,41^ using two independent simulations (total of 4 μs per complex) of RBD-ACE2 complexes for both viruses. We used two ACE2 helices (residues 21 to 83) for alignment because both helices are part of the RBD-ACE2 interface. We found that three leading modes capture more than 60% of the fluctuations of the system (Figs. 2a,b). Remarkably, all three modes are quite similar in both viruses, with the exception of mode #2, which reveals the motion in the SARS-CoV loop (residues 458-475) that transiently loses contact with the ACE2 receptor (Figs. 2c,d). Overall, normal mode #2 in SARS-CoV represents the largest movement of the RBD away from the ACE2. Similar normal modes (modes #2 and #3) were obtained from a principal component analysis in a prior study using 200-ns MD simulations.^24^

### Identification of residues distinguishing SARS-CoV from SARS-CoV-2 through machine learning

Our MD simulations showed subtle, yet important differences between the SARS-CoV and SARS-CoV-2 RBD interactions with ACE2. However, long MD simulations generate a large amount of high-dimensional data, making it difficult to identify which region or residues of the RBD make significant contributions to these differences.^33,34,36^ ML approaches can be excellent tools for identifying differences in MD trajectories^34,35,37,42^ and have recently been applied in computational studies related to SARS-CoV-2.^35,43,44^ Inspired by the work of Fleetwood et al.,^34^ we trained ML classifiers to distinguish between the configurations from the SARS-CoV and SARS-CoV-2 trajectories, and, in the process, rate the importance of each feature to the classification. The feature importance profile, hence, reveals specific residues that exemplify the different behaviors of the two RBDs.

For input features, we used inverse contact distances between residues in the RBD and in ACE2, emphasizing the differences between interacting residues across the interface. A residue pair was only included if its distance ever fell below 15 Å, resulting in a dataset of 4886 features. The distance of 15 Å is large enough to capture all contacts between the RBD and ACE2 and also to provide a reasonable speed for ML training. To minimize the bias from a particular model and increase robustness of the results, three different architectures of supervised ML classifiers were used: a linear logistic regression (LR) model, a tree-based random forest (RF) model, and a multilayer perceptron (MLP) neural network.

The three classifiers were trained to distinguish configurations of SARS-CoV RBD bound to ACE2 from those of SARS-CoV-2 RBD bound to ACE2. Highly correlated features were deleted before training. Different choices of removal generate different input data sets, which leads to different importance profiles. The choices of deletion were shuffled multiple times and the resulting importance profiles averaged until they were found to be converged (Fig. S5). The feature importance profiles were extracted from three classifiers, allowing us to focus on distinct RBD residues. The results from each model were compared against each other for cross-validation (Figs. 3, S6).

**Figure 3.**
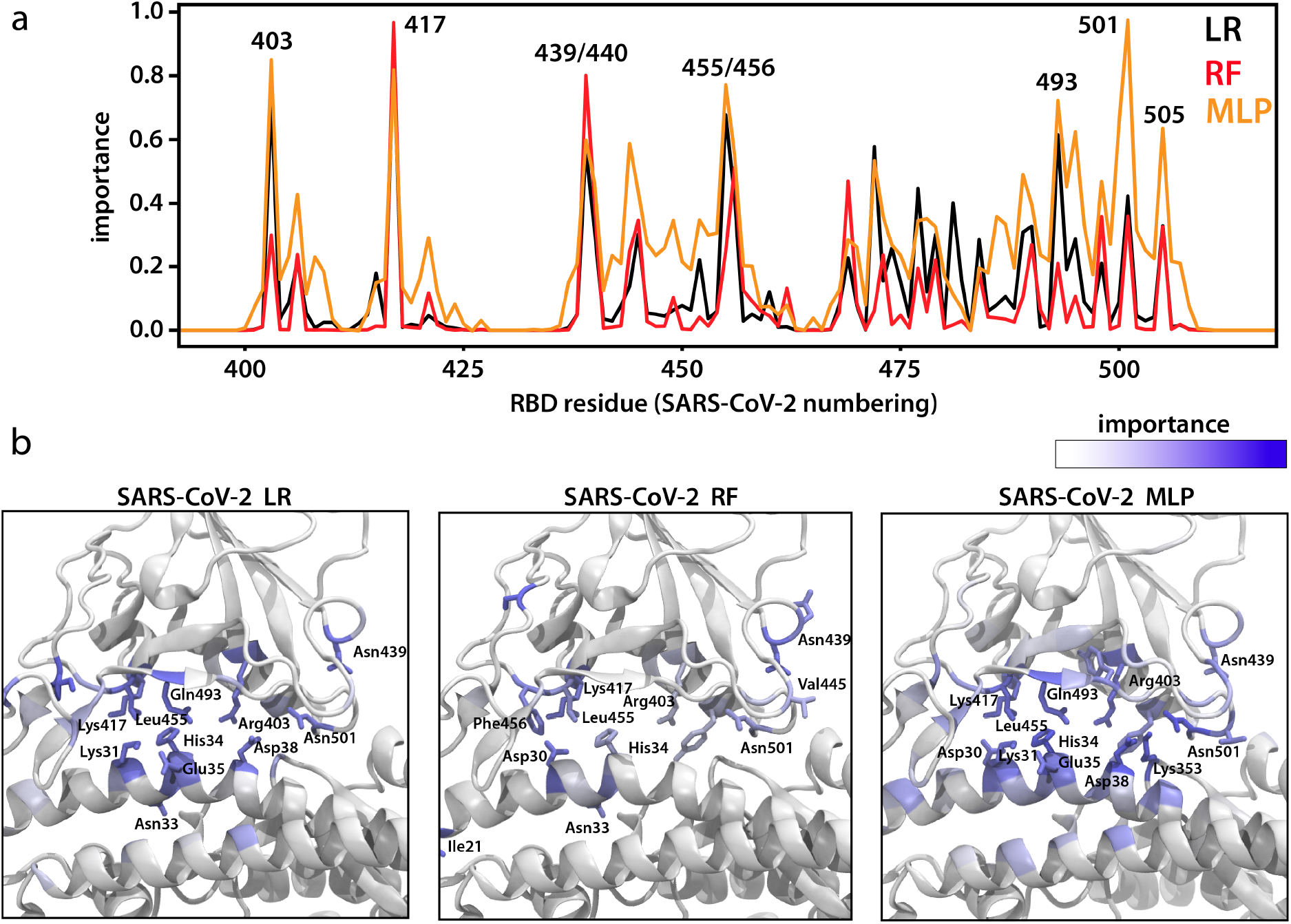
Residues determined to be most important for distinguishing between SARS-CoV and SARS-CoV-2 binding to ACE2. (a) Logistic regression (LR), random forest (RF), and multilayer perceptron (MLP) were used to identify residues that contribute the most to differences between the two RBD-ACE2 complexes. A residue with a high importance is composed of highly distinguishing distance pair(s) between RBD and ACE2 proteins in the two systems. (b) Important residues are highlighted in the SARS-CoV-2 system for each ML approach.

Overall, the per-residue importance profiles from the three methods are generally similar, sharing many of the same major peaks. In particular, Arg403, Lys417, Asn439, Leu455, and Asn501 (SARS-CoV-2 numbering) were highlighted with importance larger than 0.7 by at least one approach. It is worth noting that although the ML classifiers were trained without any knowledge of which residues are different between SARS-CoV-2 and SARS-CoV, the peak residues identified are almost all mutated between the two viruses. In addition, a cluster of minor peaks were observed for the loop containing residues 471-488. Coloring the structure by the importance values (Fig. 3b), it becomes apparent that these residues represent a particular region on the RBD that interacts with the N-terminus of ACE2. From the trajectories, we observed momentary, albeit repetitive detachment of the loop containing residues 458-475 from ACE2 in the SARS-CoV simulations, while the corresponding region maintained stable contacts with ACE2 in the SARS-CoV-2 simulations. Such detachment was accompanied by a tilting motion of the whole SARS-CoV RBD, as also captured in mode #2 of the EDA (Figs. 2c,d).

### Free-energy perturbation of ML-identified residues

While the preceding analysis ranks the relative importance of specific residues to discriminate between the conformations of SARS-CoV and SARS-CoV-2 RBDs bound to ACE2, it does not quantify the differences. FEP calculations, on the other hand, provide a means by which the energetic contribution of each mutated residue to ACE2 binding can be quantified precisely. We selected residues ranked highly by ML approaches, namely Arg403, Lys417, Asn439, Leu455, Phe456, Gln498, and Asn501 (all SARS-CoV-2 numbering) for mutation to their SARS-CoV counterpart (Table 1). Finally, the differences in interactions of these residues with ACE2 were examined in our 2 × 2-μs-long simulations of the RBD from each virus.

**Table 1.** Calculated free energy changes for mutations predicted to be important by ML. The residues in the SARS-COV-2 RBD were mutated to the corresponding SARS-CoV residues. A positive ΔΔ*G* indicates that the mutation is unfavorable for binding to ACE2.

| mutation | $\Delta\Delta G$ (kcal/mol) |
| --- | --- |
| R403K | $+1.0 \pm 1.0$ |
| K417V | $+2.2 \pm 0.9$ |
| N439R | $-3.0 \pm 0.7$ |
| L455Y | $+3.5 \pm 0.9$ |
| F456L | $+0.6 \pm 1.0$ |
| Q498Y | $-4.0 \pm 0.5$ |
| N501T | $-0.7 \pm 0.9$ |

The top-ranked residue in both RF and LR approaches (and highlighted in MLP as well) is Lys417 (Val404 in SARS-CoV). In the complex between SARS-CoV-2 and ACE2, Lys417 forms a salt bridge with Asp30 of ACE2 (Fig. 4b). Mutation to valine eliminates this salt bridge, and, from FEP, we find that Lys417 is 2.2 kcal/mol more favorable than valine at the same position. This contribution is in quantitative agreement with previous calculations.^24^

**Figure 4.**
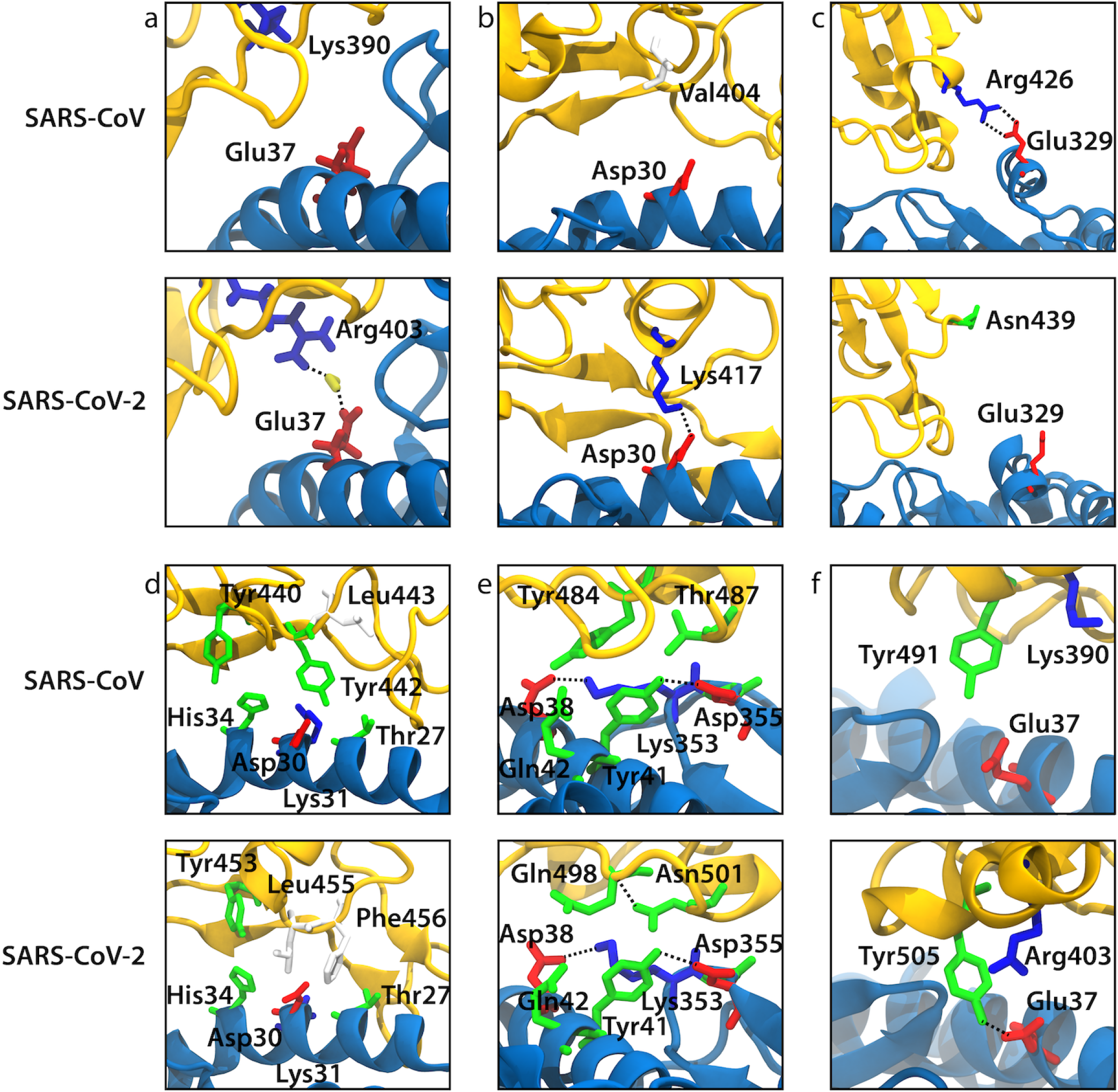
Snapshots from our MD simulations that illustrate the difference in ACE2 binding between SARS-CoV and SARS-CoV-2 for the selected mutations. ACE2 and RBD backbones are shown in blue and yellow, respectively. (a) Arg403/Lys390 (SARS-CoV-2/SARS-CoV numbering). (b) Lys417/Val404. (c) Asn439/Arg426. (d) Leu455/Tyr442. (e) Gln498-Asn501/Tyr484-Thr487. (f) Tyr491/Tyr505.

The addition of a salt bridge between ACE2 and the RBD in SARS-CoV-2 is balanced by the loss of another, namely Arg426 in SARS-CoV (Asn439 in SARS-CoV-2) with Glu329 on ACE2 (Fig. 4c). The presence of a salt bridge here stabilizes a single helical turn (Arg426 to Asp429) in the SARS-CoV RBD that is absent in SARS-CoV-2 RBD. From FEP, we find that Asn439 is 3.0 kcal/mol less favorable than arginine at the same position.

It was unexpected that Arg403 (Lys390 in SARS-CoV) would present a notable difference, given the conservation of a positively charged residue between the two RBDs. However, it was the second-ranked residue according to the LR and MLP approaches (tenth-ranked according to the RF approach). Both residues form a salt bridge with another RBD residue. The slightly bulkier Arg403 interacts with Asp405, whereas Lys390 interacts with Glu406 (SARS-CoV-2 numbering). A close inspection of the trajectories reveals that while neither Arg403 in SARS-CoV-2 nor Lys390 in SARS-CoV directly interacts with ACE2, the former comes closer, due to its interaction with Asp405. In particular, we often observed Arg403 forming a water-bridged interaction with Asp37 in ACE2, i.e., both residues formed a hydrogen bond with the same water molecule (Fig. 4a). This water-bridged interaction was observed 29% of the time. FEP calculations demonstrate that Arg403 is slightly more favorable (by 1.0 kcal/mol) at this position than lysine.

We also found that Leu455 is significantly more favorable (by 3.5 kcal/mol) than its counterpart in SARS-CoV, Tyr442. Mutation of Leu455 to tyrosine disrupts the salt bridge between Lys417 in the RBD with Asp30 in ACE2. Although this salt bridge is not present in SARS-CoV, Tyr442 forms a hydrogen bond with Asp30 instead roughly half of the time in simulations of the complex between the SARS-CoV RBD and ACE2. Next to Leu455, Phe456 makes significant hydrophobic contact with Thr27 on ACE2 (average contact area of 44.1Å^2^), as recognized previously^39^; the corresponding SARS-CoV residue, Leu443, does not (average contact area of 18.0Å^2^). Phe456 is 0.6 kcal/mol more favorable than leucine at the same position. In agreement, experiments also demonstrate that both the L455Y point mutation and the F456L point mutation significantly diminish binding of the SARS-CoV-2 RBD to ACE2.^20^ The three residues Tyr453, Leu455, and Phe456 are also more compact than their SARS-CoV counterparts (Tyr440, Tyr442, and Leu443; Fig. 4d), which allows for more stable interactions with His34 on ACE2.

Residues Gln498 and Asn501 (Tyr484 and Thr487 in SARS-CoV) are near one end of the RBM. FEP indicates that the SARS-CoV residues are more favorable in both positions, by 4.0 kcal/mol (Q498Y) and 0.7 kcal/mol (N501T). Experiments have also demonstrated the favorability of tyrosine and threonine in these two positions, respectively.^20^ The large difference for Gln498 compared to tyrosine in the same position appears to arise in part from a hydrogen bond that forms between Tyr484 and Asp38 in the SARS-CoV simulations, whereas in the SARS-CoV-2 simulations, the equivalent Gln498 forms hydrogen bonds less frequently with Asp38, Tyr41, Gln42, and Lys353. In total, Tyr484 in the SARS-CoV RBD forms hydrogen bonds with ACE2 42% of the time, whereas Gln498 in the SARS-CoV-2 RBD only forms hydrogen bonds 20% of the time. The differences between Asn501 (SARS-CoV-2) and Thr487 (SARS-CoV) are less apparent. The carbonyl on the side chain of Asn501 forms a persistent hydrogen bond with the backbone of Gln498; the side chain of the equivalent Thr487 in SARS-CoV is too short to form this hydrogen bond (Fig. 4e). The nitrogen on the side chain of Asn501 then points towards the oxygen on the side chain of Tyr41 on ACE2, but it is unable to form a hydrogen bond due to an unfavorable angle.

We also examined Tyr505 (Tyr491 in SARS-CoV), which is the same in both viruses, yet has high importance according to the MLP approach. This residue has increased hydrogen bonding with Glu37 in SARS-CoV-2 in comparison to SARS-CoV; a hydrogen bond was present 54% and 15% of the time, respectively. Upon further inspection of the simulation trajectories, we observed a cation-π interaction between Arg403 and Tyr505 that stabilizes the position of the latter; Lys390 in SARS-CoV does not form a similar interaction with Tyr491 (Fig. 4f), possibly due to the lack of dispersion contributions in lysine-arene interactions.^45^ Arg403 was one of the most important residues selected by ML for discriminating between the two viral RBDs (Fig. 3), and this importance extends to Tyr505 as well. Deep mutational scanning reveals that tryptophan is more favorable at this position, while phenylalanine and histidine are only slightly unfavorable (all others are very unfavorable),^46^ further supporting the role of a cation-π interaction in stabilizing this residue.

### Free-energy perturbation of RBD variants

Throughout the course of the pandemic, multiple variants of SARS-CoV-2 have emerged and have been identified. While some of these variants have rapidly grown in frequency, several others identified early in the pandemic still have low frequencies. Such is the case, for example, for G476S, V483A, and V367F.^1,5,47^ As above, we carried out FEP calculations for each of these three mutations, finding either very little or slightly unfavorable impact on binding to ACE2 (−0.2 to +0.4 kcal/mol; Table 2). Their small impact is commensurate with their low prevalence in the population overall.

**Table 2.** Effect of point mutations on binding to ACE2 for selected residues found in virus variants.

| mutant | $\Delta\Delta G$ (kcal/mol) |
| --- | --- |
| V367F | +0.1 ± 0.9 |
| G476S | +0.4 ± 0.4 |
| V483A | -0.2 ± 0.1 |
| K417N | +0.4 ± 0.6 |
| E484K | -1.3 ± 0.2 |
| N501Y | -4.5 ± 0.7 |

In December 2020, new variants from South Africa (lineage B.1.351) and the United Kingdom (lineage B.1.1.7) were discovered. Among a number of mutations, one common to both is N501Y, which has now also been observed in a Brazilian variant (lineage P.1).^6^ Notably, deep mutational scanning also showed that threonine, tyrosine, tryptophan, and phenylalanine are all more favorable for binding to ACE2 than Asn501 at the same position,^46^ presaging the emergence of this mutation. Supporting this conclusion, our FEP calculation shows a large decrease in binding free energy of −4.5 kcal/mol for N501Y (Table 2). This decrease is due to multiple interactions, including the formation of a hydrogen bond between Tyr501 and Lys353 on ACE2, which was also observed in another recent simulation study (Fig. S7c)^48^, hydrophobic contacts between the aliphatic portion of the Lys353 side chain and the aromatic ring of Tyr501, and a T-shaped *π*-*π* interaction of Tyr501 with Tyr41.

We also examined the E484K mutation, which was originally identified in the South African variant 501Y.V2 and was later found in the Brazilian variant 501Y.V3. More recently, this mutation has also been observed in some U.K. variants. Our FEP calculation also shows a decrease in binding free energy of −1.3 kcal/mol for E484K (Table 2). This appears to be due to a favorable electrostatic interaction between Lys484 and Glu75 on ACE2 (Fig. S7b). However, they only transiently come close enough to form a salt bridge, apparently limited by the rigidity of the loop containing Lys484, which is stabilized by a disulfide bond between Cys480 and Cys488. The weaker effect of E484K compared to N501Y is also borne out by experimental binding assays in which N501Y improved affinity for ACE2 by a factor of 4, whereas E484K only enhanced binding by a factor of 1.5.^49^

Finally, we considered the K417N mutation, found in the South African and Brazilian variants. This mutation eliminates a salt bridge between Lys417 and Asp30 (see Fig. 4b). Unsurprisingly, our FEP calculation shows that it is slightly detrimental to binding, with Asn417 increasing the free energy by 0.4 kcal/mol, in agreement with experimental binding assays.^49,50^ However, K417N has been found in simulations to have an even more detrimental effect on monoclonal antibody binding.^51^

## Discussion

The increased circulation of SARS-CoV-2 compared to SARS-CoV and MERS-CoV has been attributed, in part, to increased infectivity,^1^ for which the affinity of the viral spike protein to the target receptor plays a central role.^8,22,52^ Recent mutations of SARS-CoV-2 have increased infectivity, and many of them are believed to facilitate binding to ACE2.^5,53–55^ Therefore, it is imperative to understand the mechanisms behind enhanced binding to ACE2 by SARS-CoV-2 RBD and how binding changes due to emerging mutations, notably K417N, E484K and N501Y.^6,49^ Here, we have used a combination of long MD simulations to study the differences in dynamics between the RBDs of SARS-CoV and SARS-CoV-2 in complex with human ACE2, ML to identify the important residues, and FEP calculations to quantify the contributions of these residues to binding ACE2. This combination of computational approaches revealed key structural and energetic differences in binding to ACE2 for the two viruses (Fig. 4).

Our initial analysis of MD simulations showed minor differences in the dynamics between the RBD-ACE2 complexes for the two viruses, similar to previous, shorter MD simulations.^21,24,39^ Decreased RMSDs were observed for the SARS-CoV-2 RBD-ACE2 complex compared to SARS-CoV. Essential dynamics analysis showed that the dynamics of the two complexes are highly similar, with most of the differences occurring in the loop containing residues 471-488 (SARS-CoV-2 numbering), which was also found to dissociate temporarily from ACE2 in one of our SARS-CoV simulations.

To investigate the impact of individual residues on binding, we used three ML approaches and analyzed inverse contact distances between residues in RBD and ACE2 in the two complexes. With these approaches, we identified key residues that are responsible for differences in receptor binding. Although these three ML approaches have very different architectures, namely a linear model (LR), an ensemble of trees (RF), and a neural network (MLP), they highlight many of same residues (Arg403, Lys417, Asn439, Leu455) and also capture the motion of loop residues 471-488 (Fig. 3a). Almost all high-importance residues were mutated in SARS-CoV-2 compared to SARS-CoV. Six of the seven mutations identified in our ML approaches were also highlighted in previous studies: Val404/Lys417, Arg426/Asn439, Tyr442/Leu455, Leu443/Phe456, Tyr484/Gln498, Thr487/Asn501,^13,14,23,24^ demonstrating the ability of our approach to replicate human insight.

A previous study using MLP to identify important residues between different protein configurations showed that more hidden layers led to lower accuracy.^34^ Here, we reached an accuracy of 1 for classifying SARS-CoV from SARS-CoV-2 with only one hidden layer in the MLP implementation. If we remove all hidden layers of an MLP model, it will have the same architecture as a linear model, such as the LR approach also used here, for which the accuracy was 1 as well. Hidden layers in a neural network construct connections between features, e.g., between individual pixels in image recognition, which by themselves hold very little identifying information. However, in the RBD-ACE2 complex simulations, a single distance between two residues (one feature) can sometimes be sufficient on its own to discriminate between them, such as a distance pair including SARS-CoV-2 RBD Lys417. Instead of improving the separation between the two RBDs, the hidden layer in MLP appears to have led to a more dispersed importance profile by creating abstract connections between features (Fig. 3b). While the benefits of these connections appear limited in our application, the MLP approach did highlight one conserved residue, Tyr491/Tyr505, compared to the other two approaches. This residue was found to adopt distinct conformations in the two RBDs due to the influence of neighboring Lys390/Arg403, which itself was determined to be important by LR and MLP but not RF.

Our FEP results are in agreement with deep mutational scanning experiments by Star et al.^46^ as well as other experimental assays, demonstrating that our approach is a fast, accurate way to predict the effect of emerging SARS-CoV-2 mutations in the RBD. As in previous computational and experimental studies, although most mutations from SARS-CoV to SARS-CoV-2 were favorable, there were several exceptions, notably N439R, Q498Y, and N501T. As expected, these mutations decreased interactions with ACE2 (Fig. 4). Some of these residues have been further mutated in new variants of SARS-CoV-2, e.g., N501Y. Significant differences in free energies (>1 kcal/mol) were observed for five of the seven mutations selected by our ML approaches, indicating that they accurately identified the most meaningful mutations for subsequent FEP calculations.

Finally, we have used FEP to study the effects of several emerging mutations in SARS-CoV-2 RBD. As expected, several mutations that were observed initially, yet never became widespread, namely G476S, V483A, and V367F,^1,47^ had negligible or slightly detrimental effects on binding to ACE2 (Table 2). In contrast, two of the widespread mutations found in several strains, N501Y and E484K, greatly enhanced binding to ACE2, by −4.5 and −1.3 kcal/mol, respectively. Although the K417N mutation was unfavorable for ACE2 binding, the effect was small (+0.4 kcal/mol), and could be compensated by decreased antibody binding.^49,51^ Our results suggest that the effects on ACE2 binding play a central role in the natural selection of mutations in SARS-CoV-2 RBD, and that mutations with strongly detrimental effects on ACE2 binding are unlikely to become widespread. Our work demonstrates the utility of ML approaches for analyzing long MD trajectories, which could be generalized readily to other complexes.

## Methods

### System building

The SARS-CoV RBD bound to ACE2 was initialized from chains E and A, respectively, of the crystal structure in PDB ID: 2AJF.^56^ Missing residues 376-381 in a loop of the RBD were modeled and added. Disulfide bonds between ACE2 residues Cys133 and Cys141, Cys344 and Cys361, and Cys530 and Cys542 were added; they were also added for RBD residues Cys323 and Cys348, Cys366 and Cys419, and Cys467 and Cys474. Based on its local electrostatic environment, Asp350 of ACE2 was protonated. Crystallized glycans bound to ACE2 residues Asn90, Asn322, and Asn546, as well as RBD residue Asn330, were included.

For the SARS-CoV-2 RBD bound to ACE2, a refined version of chains E and B, respectively, from the cryo-EM structure in PDB 6M17 was used.^12,57^ In addition to the disulfide bonds for ACE2 listed above, we also added them for RBD residues Cys336 and Cys361, Cys379 and Cys432, and Cys480 and Cys488. Asp350 of ACE2 was protonated just as in the complex with SARS-CoV RBD. More bound glycans were resolved in this structure than in the SARS-CoV RBD complex. Specifically, we included those bound to ACE2 residues Asn53, Asn90, Asn103, Asn322, Asn432, and Asn546 in addition to those bound to RBD residue Asn343.

ACE2 contains a Zn^2+^ ion bound to His374, His378, Glu402, and a water molecule in a tetrahedral arrangement. The initial bonded parameters and charges for the zinc complex were taken from the ZAFF force field.^58^ Only the charges of the zinc ion and the bonded nitrogen and oxygen atoms were taken from ZAFF. The charges of carbons adjacent to these nitrogen and oxygen atoms were adjusted to maintain a total complex charge of +1. All other atoms retained their CHARMM36m charges. Only bonded parameters containing the zinc ion were taken from ZAFF, while CHARMM36m parameters were used for everything else. The initial bonded ZAFF parameters caused deformation of the tetrahedral geometry around the metal center. Therefore, parameters for the angles centered on the zinc atom were adjusted. The equilibrium values were set to 109.5°, corresponding to a perfect tetrahedral arrangement, and the force constant was increased to 130.0 kcal·mol^−1^ · rad^−2^ in order to increase the likelihood of tetrahedral geometry. It should be noted that the zinc complex is located far away from the ACE2-RBD interface and is not directly involved in binding.

Crystallographic water molecules were retained. After adding all missing hydrogen atoms, each of the two systems, SARS-CoV RBD bound to ACE2 and SARS-CoV-2 RBD bound to ACE2, was solvated in a box containing ~64,000 water molecules and ionized with Na^+^ and Cl^−^ ions to a salt concentration of 150 mM. Final system sizes were ~200,000 atoms and ~(130Å)^3^.

### MD simulations

All simulations used the CHARMM36m force field for proteins,^59^ CHARMM force field for glycans,^60^ and TIP3P water.^61^ Each system was equilibrated in multiple stages using NAMD 2.13.^62^ In the first stage, the proteins and glycans were restrained for 1 ns while water and ions were equilibrated. In the second stage, only the protein backbones were restrained to their starting positions and the side chains and glycans were equilibrated for 4 ns. Finally, all restraints were released for 10 ns. NAMD simulations were run at constant temperature (310 K) and constant isotropic pressure (1 atm) enforced by a Langevin thermostat and piston, respectively. A 2-fs time step was used, with long-range electrostatics calculated every other time step using the particle-mesh Ewald method.^63^ A short-range cutoff for Lennard-Jones interactions was set at 12 Å, with a switching function beginning at 10 Å.

After equilibration in NAMD, each of the two systems was run in duplicate for 2 μs using Amber16 on GPUs,^31^ giving 8 μs of simulation in total. A uniform 4-fs time step was employed through the use of hydrogen mass repartitioning.^64,65^ Other simulation parameters were identical to those used in the NAMD simulations with the exception of pressure control, which used a Monte Carlo barostat.

### Free-energy perturbation

The relative ACE2-RBD binding free energy associated with the mutation of selected amino acids of the SARS-CoV RBD was determined using free-energy perturbation (FEP).^66,67^ Toward this end, the amino-acid replacements were carried out in the free RBD (unbound state) on the one hand, and in the ACE2-RBD complex (bound state) on the other hand. Considering the nature of the alchemical transformations at hand, the reaction path was stratified into 100 windows of equal widths. Replacement of the WT residue by an alternate one was performed over 160 ns, both in the bound and unbound states. The dual-topology paradigm was used,^68^ whereby the initial and the final states of each alchemical transformation coexist but do not interact. Use was made of the library of amino-acid hybrids compliant with the CHARMM36m force field for proteins,^59^ available in the visualization program VMD.^69^ To augment sampling efficiency wherever necessary, geometric restraints were introduced to ensure that the *β*– and γ–carbon atoms of the alternate topologies remained superimposed during the course of the amino-acid replacement. In each window of the reaction path, data collection was prefaced by suitable thermalization in the amount of one fourth of the total sampling. To improve the accuracy of the different free-energy calculations, each alchemical transformation was carried out bidirectionally,^70^ and the associated relative binding free energy was computed using the Bennett acceptance ratio (BAR) method.^71^ The error bars associated with each relative binding free energy were determined on the basis of the hysteresis between the forward and backward alchemical transformations of the bidirectional FEP calculations, assuming independence of the transformations in the bound and unbound states. All free-energy calculations were performed with the single-node path implementation of FEP^72^ available in NAMD 3.0.^32^

### Machine learning

The contact distances, defined as the closest distance between heavy atoms of residues on the ACE2 and RBD region of spike protein were extracted from the simulation trajectories. Only residue pairs with distances less than 15 Å in at least one frame were used as features for machine learning, forming a dataset of 4886 features. The distances were inverted and normalized to have a mean of 0 and standard deviation of 1. Highly correlated features were removed with a threshold of 0.9. Different choices of removal led to different datasets and importance profiles. Converged results were derived by averaging the importance profiles of different datasets generated by shuffling the choices 20 times (Fig. S5). Because all machine learning methods used here are supervised learning, all data points were labeled as CoV or CoV-2. We tuned the hyperparameters for each ML approach to reach an accuracy of 1, i.e., all frames were correctly classified as from SARS-CoV or SARS-CoV-2 simulations. The residue importance was derived by summing the importance of distance pairs containing a given residue and normalizing the results to have a maximum of 1.

### Logistic Regression

Logistic regression (LR) is similar to linear regression but uses a sigmoidal function,

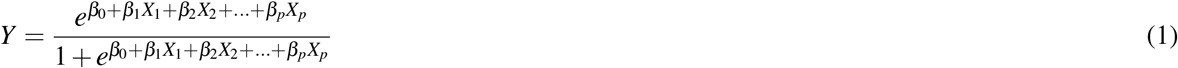

where *X_i_* is an inputted feature and *Y* is the output whose range is between 0 and 1. In classification, the data points are classified according to an output threshold value of 0.5. The LR implementation in the scikit-learn library was used.^73^ The importance of a feature is the corresponding weight (*β_1_* to *β_p_*, normalized) in equation (1). When dividing data into training and test sets, different random states led to different importance profiles in the LR classifier (Fig. S3a). Bootstrapping was used to generate multiple training sets and to obtain converged average importance profiles in the LR classifier (Fig. S3b).

### Random Forest

A random forest (RF) is composed of multiple decision trees. The classification result of one random forest is decided by the majority vote from all of the decision trees. During the training process, the decision tree algorithm iteratively searches features and determines a threshold value to split the dataset in such a way that the lowest Gini impurity as possible is obtained in each internal node. A lower Gini impurity of the split data indicates that more information is obtained. The importance of a feature is calculated as the total reduction of the Gini impurity contributed by this feature. The RF classifier has an internal bootstrapping process and generates consistent profiles (Fig. S4). We implemented RF through the scikit-learn library.^73^ We included 500 trees in our model and generated bootstrap samples when building trees. The maximum depth of each tree was 60 and the number of features considered in each node was set to 50.

### Multilayer Perceptron and Layer-Wise Relevance Propagation

A multilayer perceptron (MLP) is a type of feed-forward artificial neural network. Compared to linear classification or regression models, an MLP includes extra hidden layer(s) of perceptrons (nodes) between input and output layers. Except for the input layer, each node uses a nonlinear activation function to deal with the signals propagated from the previous layer. We used the scikit-learn implementation of MLP.^73^ We set one hidden layer with 55 nodes (roughly the square root of the product of the number of input nodes number and output nodes) using the Rectified Linear Unit (ReLU) as the activation function. ReLU was chosen because it is faster, capable of outputting true zero, and easier to optimize than sigmoid or tanh activation functions. Labels were one-hot encoded. During training, Adam,^74^ a stochastic gradient-based optimizer, was used to optimize weights between nodes. Feature importance was extracted from MLP using Layer-Wise Relevance Propagation (LRP). If MLP makes a correct prediction or classification, LRP determines which features contribute more to this decision than others. We define the relevances *R_i_* of an output layer node as 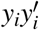, where *Y* is the true one-hot label and *Y′* is the network output. LRP propagates relevance *R* from the output layer to the input layer thought the weights of the network and neural activations. The propagation follows the LPR-0 rule,^75^ 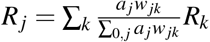, where *R_k_* and *R_j_* are the relevances of two neurons in one layer and the previous layer, separately, and *a^j^* is the activation and *w_jk_* is the weight between two neurons. The importance of one feature is calculated from the average relevance of the corresponding input neuron over all frames.

## Data Availability

## Acknowledgements

Computational resources were provided through the Extreme Science and Engineering Discovery Environment (XSEDE; TG-MCB130173), which is supported by the National Science Foundation (NSF; ACI-1548562). This work also used the Hive cluster, which is supported by the National Science Foundation under grant number 1828187 and is managed by the Partnership for an Advanced Computing Environment (PACE) at the Georgia Institute of Technology. CC acknowledges support from the Agence Nationale de la Recherche (grants ProteaseInAction and Contrats Doctoraux en Intelligence Artificielle), and from the Lorraine Université d’Excellence initiative.

## Data availability

Simulation trajectories are available at the NSF MolSSI COVID-19 Molecular Structure and Therapeutics Hub at https://covid.molssi.org.

## Author contributions statement

A.P., A.A., D.L.L., and J.C.G. designed and performed simulations. Z.Z., Y.T.P., Z.M., and J.M.P. carried out ML analysis. C.C. performed and analyzed free-energy calculations. All authors carried out analysis and wrote the manuscript.

## Notes

### Competing Interest Statement

The authors have declared no competing interest.

## References

1. Hu, B., Guo, H., Zhou, P. & Shi, Z.-L. Characteristics of SARS-CoV-2 and COVID-19. Nat. Rev. Microbiol. 1–14 (2020).

2. Xu, J. et al. Systematic comparison of two animal-to-human transmitted human coronaviruses: SARS-CoV-2 and SARS-CoV. Viruses 12, 244 (2020).

3. Rossi, G. A., Sacco, O., Mancino, E., Cristiani, L. & Midulla, F. Differences and similarities between SARS-CoV and SARS-CoV-2: spike receptor-binding domain recognition and host cell infection with support of cellular serine proteases. Infection 45, 665–669 (2020).

4. van Dorp, L. et al. Emergence of genomic diversity and recurrent mutations in SARS-CoV-2. Infect. Genet. Evol. 83, 104351 (2020).

5. Li, Q. et al. The Impact of Mutations in SARS-CoV-2 Spike on Viral Infectivity and Antigenicity. Cell 182, 1284–1294 (2020).

6. Fontanet, A. et al. SARS-CoV-2 variants and ending the COVID-19 pandemic. Lancet (2021). In press.

7. Bonilla-Aldana, D. K. et al. Bats in ecosystems and their Wide spectrum of viral infectious potential threats: SARS-CoV-2 and other emerging viruses. Int. J. Infect. Dis. 102, 87–96 (2021).

8. Li, F. Structure, function, and evolution of coronavirus spike proteins. Annu. Rev. Virol. 3, 237–261 (2016).

9. Hoffmann, M. et al. SARS-CoV-2 Cell Entry Depends on ACE2 and TMPRSS2 and Is Blocked by a Clinically Proven Protease Inhibitor. Cell 181, 271–280 (2020).

10. Shang, J. et al. Cell entry mechanisms of SARS-CoV-2. Proc. Natl Acad. Sci. USA 117, 11727–11734 (2020).

11. Li, W. et al. Receptor and viral determinants of SARS-coronavirus adaptation to human ACE2. EMBO J. 24, 1634–1643 (2005).

12. Yan, R. et al. Structural basis for the recognition of SARS-CoV-2 by full-length human ACE2. Science 367, 1444–1448 (2020).

13. Lan, J. et al. Structure of the SARS-CoV-2 spike receptor-binding domain bound to the ACE2 receptor. Nature 581, 215–220 (2020).

14. Shang, J. et al. Structural basis of receptor recognition by SARS-CoV-2. Nature 581, 221–224 (2020).

15. Wang, Q. et al. Structural and functional basis of SARS-CoV-2 entry by using human ACE2. Cell 181, 894–904.e9 (2020).

16. Hamming, I. et al. Tissue distribution of ACE2 protein, the functional receptor for SARS coronavirus. A first step in understanding SARS pathogenesis. J. Pathol. 203, 631–637 (2004).

17. Dai, L. & Gao, G. F. Viral targets for vaccines against COVID-19. Nat. Rev. Immunol. 21, 73–82 (2021).

18. Ge, J. et al. Antibody neutralization of SARS-CoV-2 through ACE2 receptor mimicry. Nat. Commun. 12, 250 (2021).

19. Robert, X. & Gouet, P. Deciphering key features in protein structures with the new ENDscript server. Nucleic Acids Res. 42, W320–324 (2014).

20. Yi, C. et al. Key residues of the receptor binding motif in the spike protein of SARS-CoV-2 that interact with ACE2 and neutralizing antibodies. Cell. Mol. Immunol. 17, 621–630 (2020).

21. Ghorbani, M., Brooks, B. R. & Klauda, J. B. Critical sequence hotspots for binding of novel coronavirus to angiotensin converter enzyme as evaluated by molecular simulations. J. Phys. Chem. B 124, 10034–10047 (2020).

22. Wu, K., Peng, G., Wilken, M., Geraghty, R. J. & Li, F. Mechanisms of host receptor adaptation by severe acute respiratory syndrome coronavirus. J. Biol. Chem. 287, 8904–8911 (2012).

23. Zou, J. et al. Computational prediction of mutational effects on SARS-CoV-2 binding by relative free energy calculations. J. Chem. Inf. Model. 60, 5794–5802 (2020).

24. Wang, Y., Liu, M. & Gao, J. Enhanced receptor binding of SARS-CoV-2 through networks of hydrogen-bonding and hydrophobic interactions. Proc. Natl Acad. Sci. USA 117, 13967–13974 (2020).

25. Delgado, J. M. et al. Molecular basis for SARS-CoV-2 spike affinity for human ACE2 receptor. bioRxiv DOI: 10.1101/2020.09.10.291757 (2020).

26. Nguyen, H. L. et al. Does SARS-CoV-2 bind to human ACE2 more strongly than does SARS-CoV? J. Phys. Chem. B 124, 7336–7347 (2020).

27. Barros, E. P. et al. The Flexibility of ACE2 in the Context of SARS-CoV-2 Infection. Biophys. J. (2020).

28. Mugnai, M. L., Templeton, C., Elber, R. & Thirumalai, D. Role of long-range allosteric communication in determining the stability and disassembly of SARS-COV-2 in complex with ACE2. bioRxiv DOI: 10.1101/2020.11.30.405340 (2020).

29. Casalino, L. et al. Beyond shielding: the roles of glycans in the SARS-CoV-2 spike protein. ACS Cent. Sci. 6, 1722–1734 (2020).

30. Zhao, P. et al. Virus-receptor interactions of glycosylated SARS-CoV-2 spike and human ACE2 receptor. Cell Host Microbe 28, 586–601 (2020).

31. Salomon-Ferrer, R., Götz, A. W., Poole, D., Le Grand, S. & Walker, R. C. Routine microsecond molecular dynamics simulations with AMBER on GPUs. 2. Explicit solvent particle mesh Ewald. J. Chem. Theory Comput. 9, 3878–3888 (2013).

32. Phillips, J. C. et al. Scalable molecular dynamics on CPU and GPU architectures with NAMD. J. Chem. Phys. 153, 044130 (2020).

33. Bernetti, M., Bertazzo, M. & Masetti, M. Data-driven molecular dynamics: A multifaceted challenge. Pharm. (Basel) 13, 253 (2020).

34. Fleetwood, O., Kasimova, M. A., Westerlund, A. M. & Delemotte, L. Molecular insights from conformational ensembles via machine learning. Biophys. J. 118, 765–780 (2020).

35. Casalino, L. et al. AI-Driven Multiscale Simulations Illuminate Mechanisms of SARS-CoV-2 Spike Dynamics. bioRxiv (2020).

36. Wang, Y., Lamim Ribeiro, J. M. & Tiwary, P. Machine learning approaches for analyzing and enhancing molecular dynamics simulations. Curr Opin. Struct. Biol. 61, 139–145 (2020).

37. Haghighatlari, M. et al. Learning to make chemical predictions: The interplay of feature representation, data, and machine learning methods. Chem 6, 1527–1542 (2020).

38. Noé, F., Tkatchenko, A., Müller, K. R. & Clementi, C. Machine Learning for Molecular Simulation. Annu. Rev. Phys. Chem. 71, 361–390 (2020).

39. Ali, A. & Vijayan, R. Dynamics of the ACE2-SARS-CoV-2/SARS-CoV spike protein interface reveal unique mechanisms. Sci. Rep. 10, 14214 (2020).

40. Amadei, A., Linssen, A. B. & Berendsen, H. J. Essential dynamics of proteins. Proteins: Struct., Funct., Bioinf. 17, 412–425 (1993).

41. Bakan, A., Meireles, L. M. & Bahar, I. ProDy: protein dynamics inferred from theory and experiments. Bioinformatics 27, 1575–1577 (2011).

42. Esposito, C., Wang, S., Lange, U. E. W., Oellien, F. & Riniker, S. Combining machine learning and molecular dynamics to predict P-glycoprotein substrates. J. Chem. Inf. Model. 60, 4730–4749 (2020).

43. Alimadadi, A. et al. Artificial intelligence and machine learning to fight COVID-19. Physiol. Genomics 52, 200–202 (2020).

44. Heo, L. & Feig, M. Modeling of severe acute respiratory syndrome coronavirus 2 (SARS-CoV-2) proteins by machine learning and physics-based refinement. bioRxiv DOI: 10.1101/2020.03.25.008904 (2020).

45. Kumar, K. et al. Cation-π interactions in protein-ligand binding: theory and data-mining reveal different roles for lysine and arginine. Chem. Sci. 9, 2655–2665 (2018).

46. Starr, T. N. et al. Deep mutational scanning of SARS-CoV-2 receptor binding domain reveals constraints on folding and ACE2 binding. Cell 182, 1295–1310.e20 (2020).

47. Chen, A. T., Altschuler, K., Zhan, S. H., Chan, Y. A. & Deverman, B. E. COVID-19 CG enables SARS-CoV-2 mutation and lineage tracking by locations and dates of interest. eLife 10, e63409 (2021).

48. Fratev, F. The N501Y and K417N mutations in the spike protein of SARS-CoV-2 alter the interactions with both hACE2 and human derived antibody: A free energy of perturbation study. bioRxiv DOI: 10.1101/2020.12.23.424283 (2021).

49. Tian, F. et al. Mutation N501Y in RBD of spike protein strengthens the interaction between COVID-19 and its receptor ACE2. bioRxiv DOI: 10.1101/2021.02.14.431117 (2021).

50. Collier, D. A. et al. Sensitivity of SARS-CoV-2 B.1.1.7 to mRNA vaccine-elicited antibodies. Nature (2021). In press.

51. Luan, B. & Huynh, T. Insights on SARS-CoV-2’s mutations for evading human antibodies: Sacrifice and survival. bioRxiv DOI: 10.1101/2021.02.06.430088 (2021).

52. Hasegawa, K. et al. Affinity thresholds for membrane fusion triggering by viral glycoproteins. J. Virol. 81, 13149–13157 (2007).

53. Thomson, E. C. et al. Circulating SARS-CoV-2 spike N439K variants maintain fitness while evading antibody-mediated immunity. Cell in press (2021).

54. Volz, E. et al. Evaluating the effects of SARS-CoV-2 spike mutation D614G on transmissibility and pathogenicity. Cell 184, 64–75.e11 (2021).

55. Wise, J. Covid-19: New coronavirus variant is identified in UK. BMJ 371, m4857 (2020).

56. Li, F., Li, W., Farzan, M. & Harrison, S. C. Structure of SARS coronavirus spike receptor-binding domain complexed with receptor. Science 309, 1864–1868 (2005).

57. Croll, T. I., Williams, C. J., Chen, V. B., Richardson, D. C. & Richardson, J. S. Improving SARS-CoV-2 structures: peer review by early coordinate release. Biophys. J. (2021). In press.

58. Peters, M. B. et al. Structural survey of zinc-containing proteins and development of the Zinc AMBER force field (ZAFF). J. Chem. Theory Comput. 6, 2935–2947 (2010).

59. Huang, J. et al. CHARMM36m: an improved force field for folded and intrinsically disordered proteins. Nat. Methods 14, 71–73 (2017).

60. Guvench, O. et al. CHARMM additive all-atom force field for carbohydrate derivatives and its utility in polysaccharide and carbohydrate-protein modeling. J. Chem. Theory Comput. 7, 3162–3180 (2011).

61. Jorgensen, W. L., Chandrasekhar, J., Madura, J. D., Impey, R. W. & Klein, M. L. Comparison of simple potential functions for simulating liquid water. J. Chem. Phys. 79, 926–935 (1983).

62. Phillips, J. C. et al. Scalable molecular dynamics with NAMD. J. Comput. Chem. 26, 1781–1802 (2005).

63. Darden, T., York, D. & Pedersen, L. Particle mesh Ewald: An *N* · log(*N*) method for Ewald sums in large systems. J. Chem. Phys. 98, 10089–10092 (1993).

64. Hopkins, C. W., Le Grand, S., Walker, R. C. & Roitberg, A. E. Long-time-step molecular dynamics through hydrogen mass repartitioning. J. Chem. Theory Comput. 11, 1864–1874 (2015).

65. Balusek, C. et al. Accelerating membrane simulations with Hydrogen Mass Repartitioning. J. Chem. Theory Comput. 15, 4673–4686 (2019).

66. Landau, L. D. Statistical physics (The Clarendon Press, Oxford, 1938).

67. Zwanzig, R. W. High-temperature equation of state by a perturbation method. I. nonpolar gases. J. Chem. Phys. 22, 1420–1426 (1954).

68. Gao, J., Kuczera, K., Tidor, B. & Karplus, M. Hidden thermodynamics of mutant proteins: A molecular dynamics analysis. Science 244, 1069–1072 (1989).

69. Humphrey, W., Dalke, A. & Schulten, K. VMD: visual molecular dynamics. J. Mol. Graph. 14, 33–38 (1996).

70. Pohorille, A., Jarzynski, C. & Chipot, C. Good practices in free-energy calculations. J. Phys. Chem. B 114, 10235–10253 (2010).

71. Bennett, C. H. Efficient estimation of free energy differences from Monte Carlo data. J. Comp. Phys. 22, 245–268 (1976).

72. Chen, H. et al. Boosting free-energy perturbation calculations with GPU-accelerated NAMD. J. Chem. Inf. Model. 60, 5301–5307 (2020).

73. Pedregosa, F. et al. Scikit-learn: Machine learning in Python. J. Mach. Learn. Res. 12, 2825–2830 (2011).

74. Kingma, D. P. & Ba, J. Adam: A method for stochastic optimization. arXiv preprint arXiv:1412.6980 (2014).

75. Bach, S. et al. On pixel-wise explanations for non-linear classifier decisions by layer-wise relevance propagation. PLOS ONE 10, e0130140 (2015).

