## Supplementary Information for "Critical interactions for SARS-CoV-2 spike protein binding to ACE2 identified by machine learning"

### Free-energy perturbation of other residues

**Table S1.** Free energy effects on binding to ACE2 for selected mutations not captured by ML and for mutations of virus variants.

| mutant | $\Delta\Delta G$ (kcal/mol) |
|---|---|
| L455F | +1.6 ± 0.4 |
| F486L | +1.0 ± 0.1 |
| Q493N | 0.0 ± 0.5 |
| Q493R | +3.8 ± 1.0 |
| Q493K | +1.3 ± 0.6 |
| S494D | +1.2 ± 0.3 |

A previous analysis compared the RBM between SARS-CoV-2 and a number of SARS-CoV variants.[1] The analysis focused on interfacial SARS-CoV-2 residues Leu455, Phe486, Gln493, Ser494, and Asn501. While Leu455 and Asn501 are described in the main text, the other three were not identified as significant by at least one of the ML approaches (Gln493 was picked out by the LR approach). For comparison, we also carried out FEP for each of the three to their SARS-CoV variant.

Phe486 in SARS-CoV-2 is equivalent to Leu472 in SARS-CoV, although intriguingly a 2008 variant of SARS-CoV has Phe472 instead.[2] Previous studies have shown that mutation to phenylalanine in this position increases hydrophobic contacts with ACE2.[3,4] Our FEP calculation also showed that phenylalanine is more favorable at this position than leucine by 1.0 kcal/mol (Table S1). Gln493 is an asparagine in SARS-CoV; FEP gave no difference between the two amino acids at this position. This position for the SARS-CoV variant that infects civets is either arginine or lysine; FEP indicates both of these are significantly worse for binding to human ACE2 (increase of 3.8 and 1.3 kcal/mol, respectively). Finally, Ser494 is an aspartate in SARS-CoV. While the latter was predicted to be better for binding due to the ability to form interactions with Lys353 on ACE2, instead we find that an aspartate here is unfavorable by 1.2 kcal/mol, likely due to nearby Glu35 and Asp38 on ACE2. Our FEP results for this mutation are in agreement with deep mutational scanning data.[5] In fact, in equilibrium simulations, we find that Asp480 on SARS-CoV almost exclusively interacts with Lys439 on the RBD instead of ACE2.

In Mugnai et al.,[6] the mutation F486L was computed via alchemical transformation to have a $\Delta\Delta G$ of +1.2±0.3 kcal/mol, and Wang et al.[4] report the inverse mutation (L472F in SARS-CoV) to have a ΔΔG of -1.2± 0.2 kcal/mol. Wang et al. also report that in SARS-CoV the replacement of the internal salt bridge, Asp480/Lys439, with the corresponding residues of SARS-CoV-2 (Ser494 and Leu452) improves binding with a $\Delta\Delta G$ of -1.9 ± 0.9 kcal/mol.

Although Leu455 was already identified as important by ML and its mutation to tyrosine from SARS-CoV examined above, it has been predicted that phenylalanine would be optimal at this position.[1] However, FEP shows that Phe455 is notably worse (1.6 kcal/mol) than Leu455. Close examination of the FEP simulations revealed that Phe455 disrupts the salt bridge between Lys417 on RBD and Asp30 on ACE2, illustrating that the effects of mutating interfacial residues spread to neighboring interactions as well.

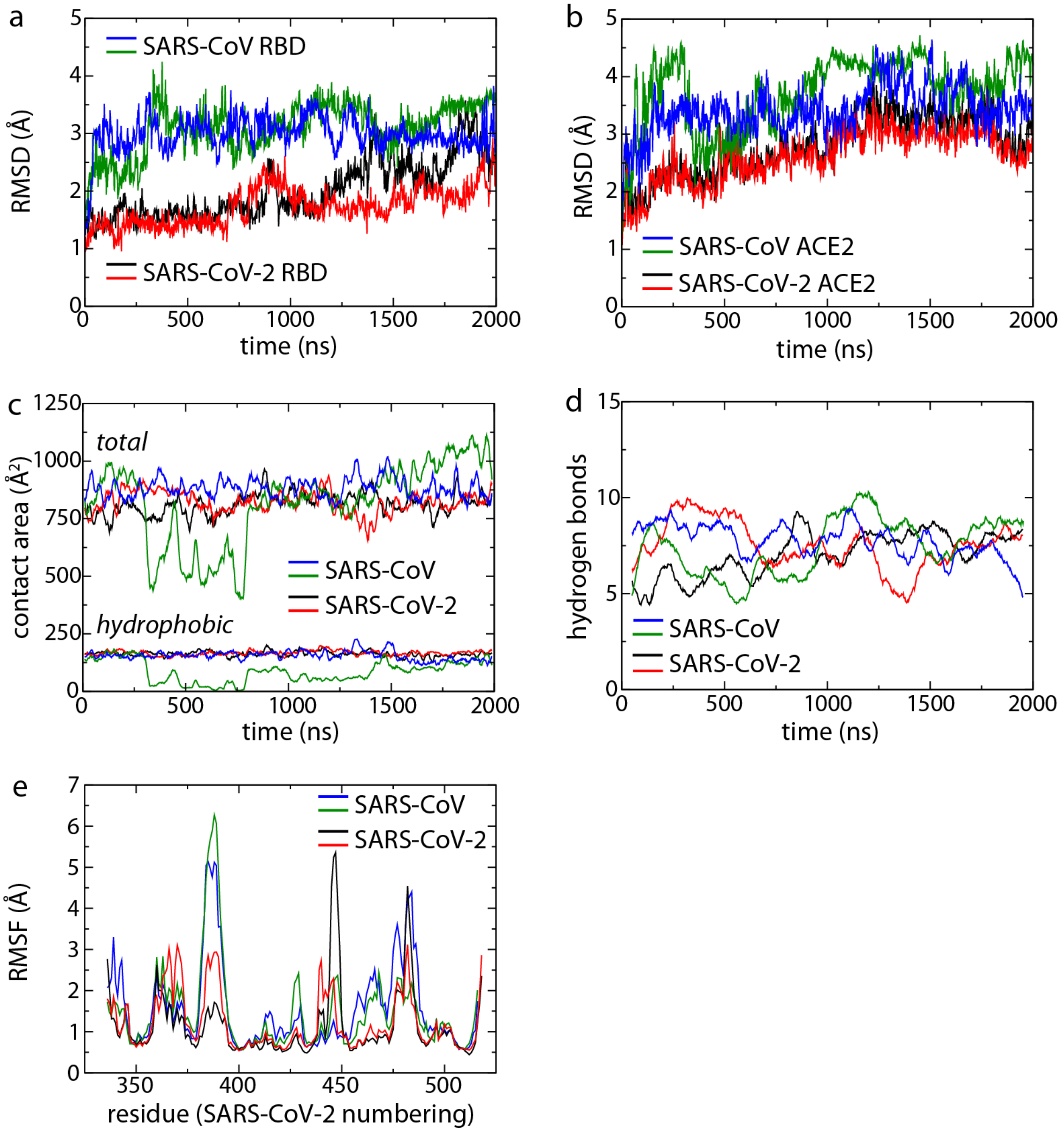


**Figure S1.** Analysis of 2-μs simulations. (a,b) Root-mean-square deviation (RMSD) of the (a) RBDs and (b) ACE2 compared to their initial structures. (c) Total and hydrophobic buried surface area between the RBD and ACE2. Area only includes that contributed by one binding partner to the interface. (d) Number of hydrogen bonds between the RBD and ACE2. Data is smoothed using a 100-ns running average. (e) Root-mean-square fluctuations (RMSF) for $C_\alpha$ atoms of each system.

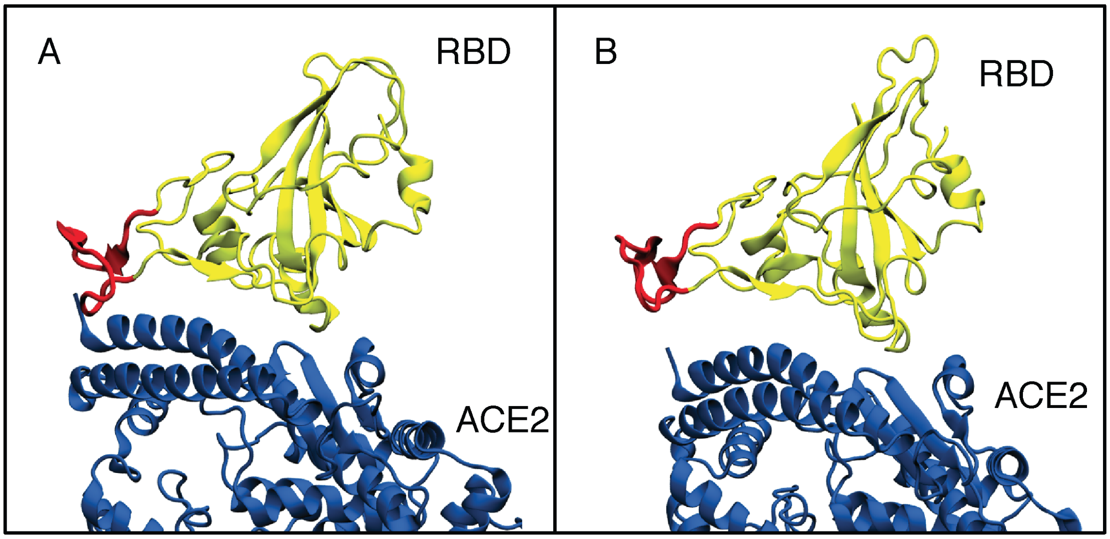


**Figure S2.** Snapshots showing movement of the loop containing the residues 458-475 (SARS-CoV numbering). The loop is shown in red, ACE2 is shown in blue, and RBD is shown in yellow. (a) The initial structure. (b) A snapshot showing the loop moving away from ACE2.

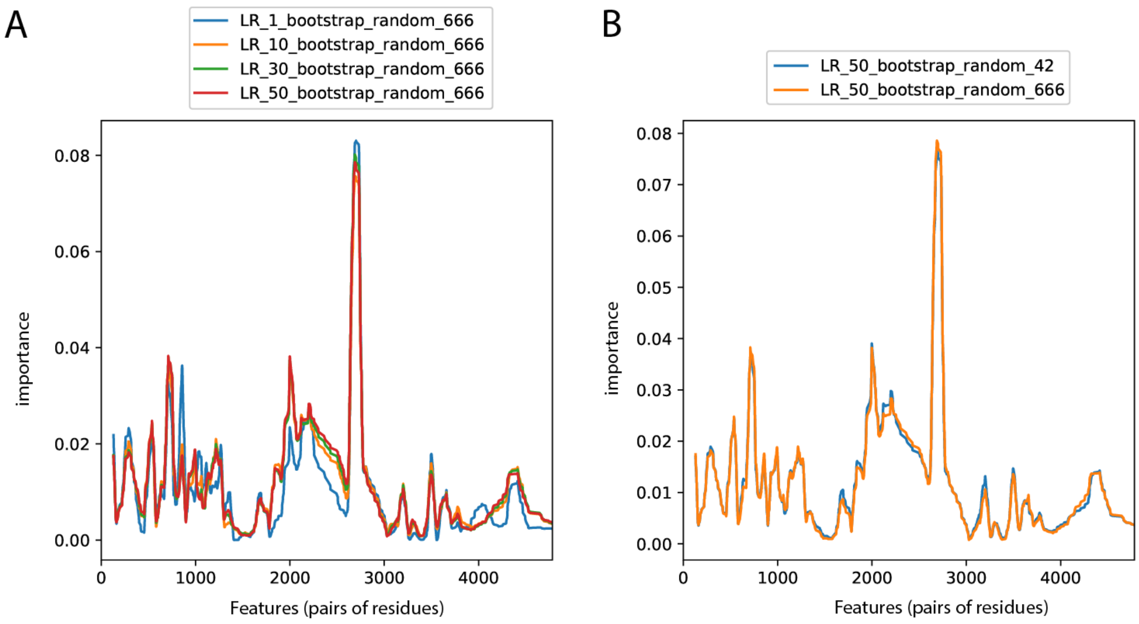


**Figure S3.** Different random states lead to different importance profiles in the LR classifier (a). Bootstrapping was used to generate multiple training sets and get converged average importance profiles for the LR classifier (b).

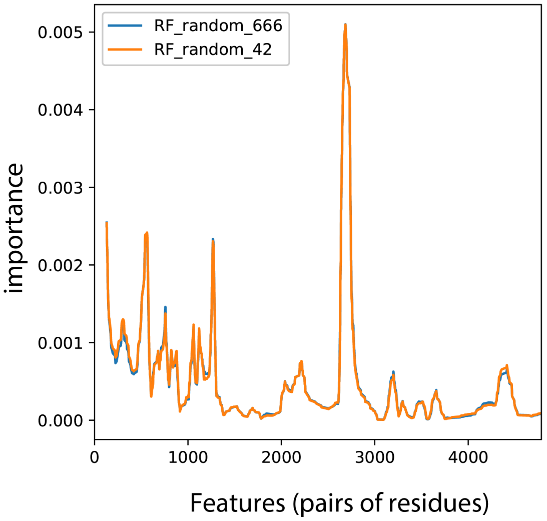


**Figure S4.** RF classifier has an internal bootstrapping process and generates consistent profiles.

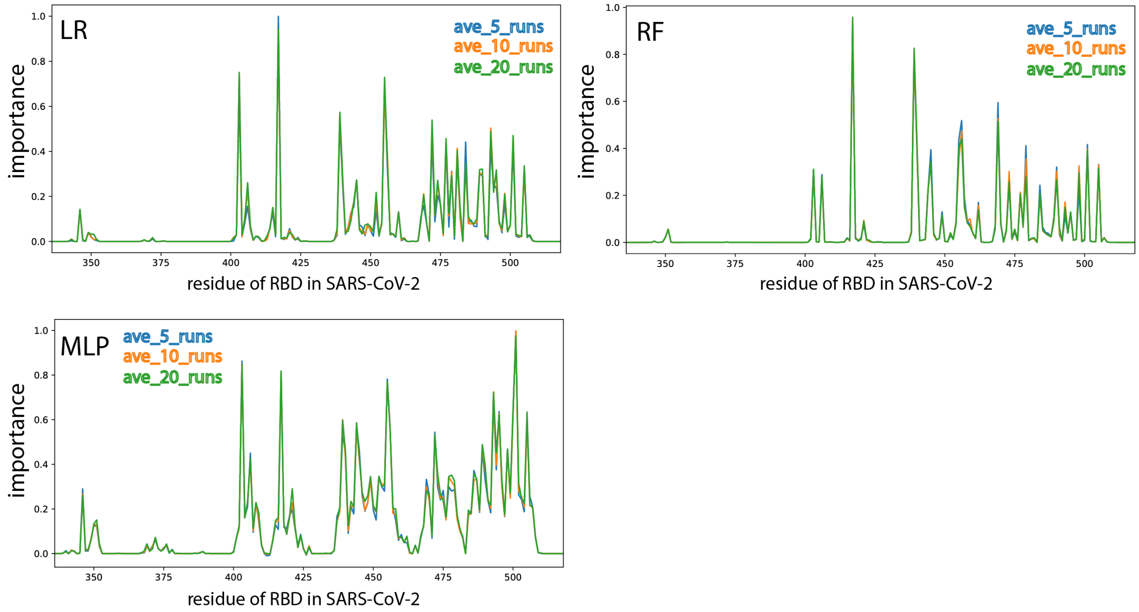


**Figure S5.** The order of deletion of highly correlated features was shuffled multiple times, and average importance profiles are converged.

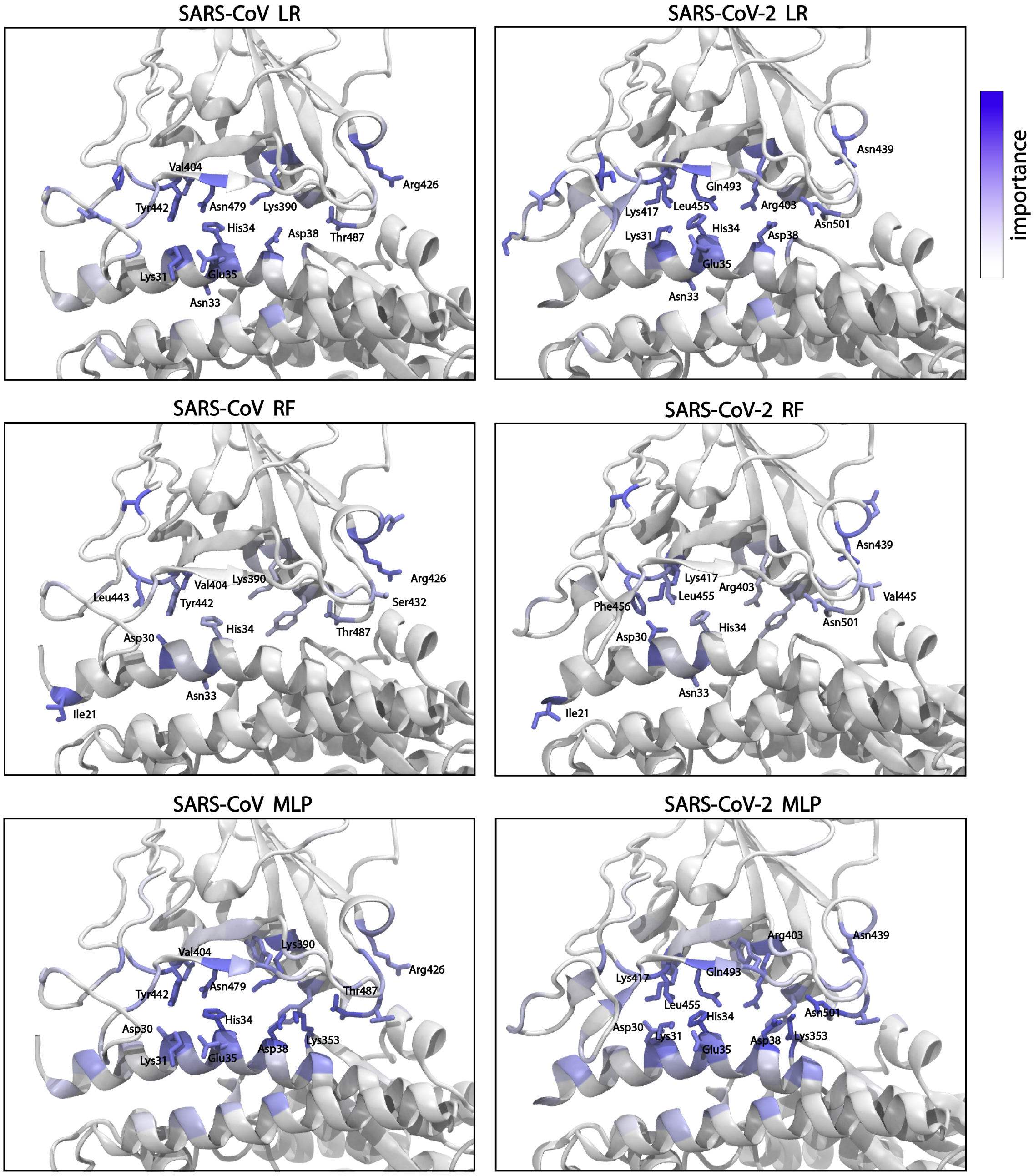


**Figure S6.** Important residues are highlighted in both SARS-CoV and SARS-CoV-2 systems with different methods (LR, RF, and MLP).

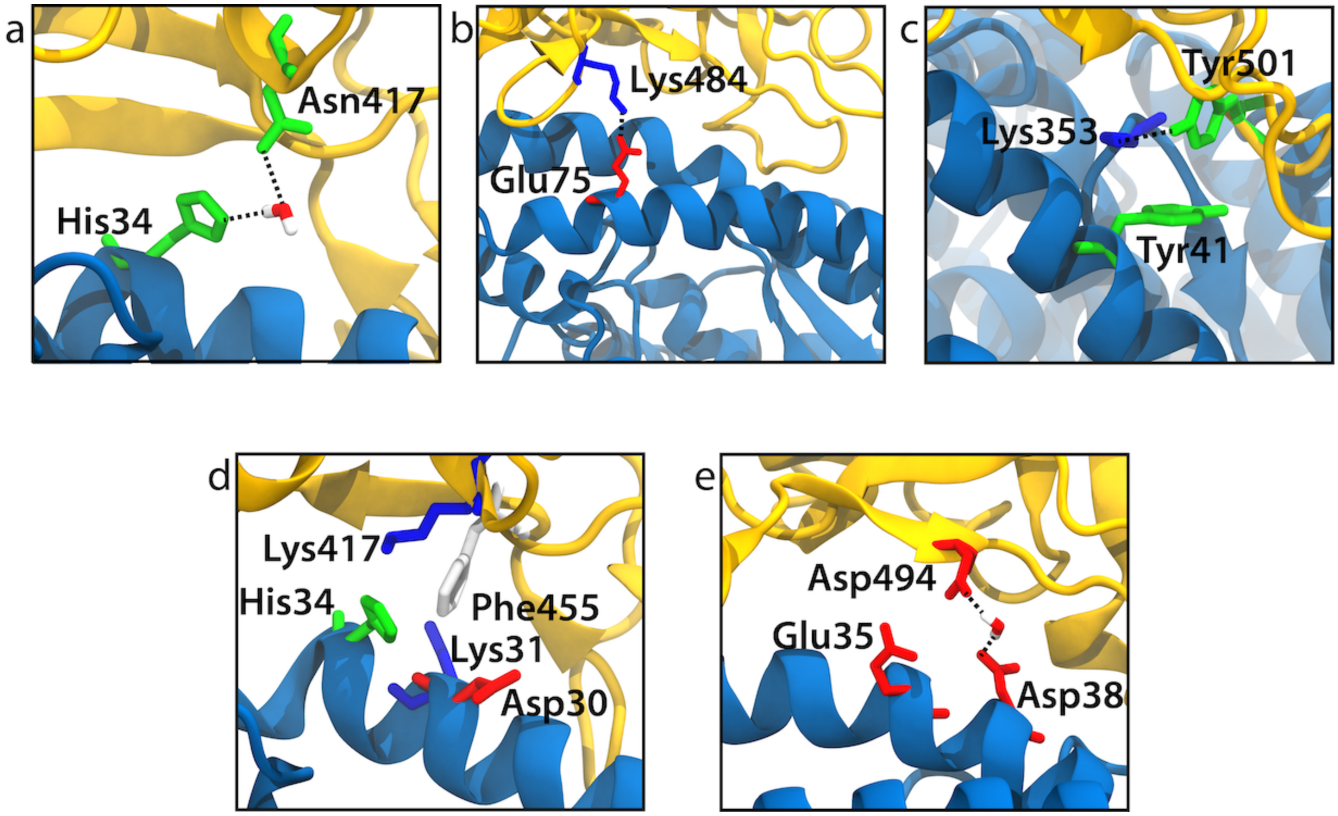


**Figure S7.** Specific interactions between the RBD (gold) and ACE2 (blue) observed in the end states of FEP calculations. (a) K417N. A water-bridged interaction between Asn417 and His34 is shown. (b) E484K. A tenuous salt bridge between Lys484 and Glu75 is shown. (c) N501Y. A hydrogen bond forms between Tyr501 and Lys353. (d) L455F. Phe455 disrupts the salt bridge between Lys417 and Asp30. (e) S494D. Despite the electrostatic repulsion between Asp494 and Glu35 and Asp38, some water-bridged interactions were observed.

### References


**1.** Wan, Y., Shang, J., Graham, R., Baric, R. S. & Li, F. Receptor Recognition by the Novel Coronavirus from Wuhan: an Analysis Based on Decade-Long Structural Studies of SARS Coronavirus. *J. Virol.* **94** (2020).

**2.** Sheahan, T. *et al.* Mechanisms of zoonotic severe acute respiratory syndrome coronavirus host range expansion in human airway epithelium. *J. Virol.* **82**, 2274–2285 (2008).

**3.** Wang, Q. *et al.* Structural and functional basis of SARS-CoV-2 entry by using human ACE2. *Cell* **181**, 894–904.e9 (2020).

**4.** Wang, Y., Liu, M. & Gao, J. Enhanced receptor binding of SARS-CoV-2 through networks of hydrogen-bonding and hydrophobic interactions. *Proc. Natl Acad. Sci. USA* **117**, 13967–13974 (2020).

**5.** Starr, T. N. *et al.* Deep mutational scanning of SARS-CoV-2 receptor binding domain reveals constraints on folding and ACE2 binding. *Cell* **182**, 1295–1310.e20 (2020).

**6.** Mugnai, M. L., Templeton, C., Elber, R. & Thirumalai, D. Role of long-range allosteric communication in determining the stability and disassembly of SARS-COV-2 in complex with ACE2. *bioRxiv* DOI: 10.1101/2020.11.30.405340 (2020).
